# Intraspecies interactions of *Streptococcus mutans* impact biofilm architecture and virulence determinants in childhood dental caries

**DOI:** 10.1101/2023.12.13.571561

**Authors:** Stephanie S. Momeni, Xixi Cao, Baotong Xie, Katherine Rainey, Noel K. Childers, Hui Wu

## Abstract

Early childhood dental caries (ECC) is the most common chronic disease among children with a heavy disease burden among low socioeconomic populations. *Streptococcus mutans* is most frequently associated with initiation of ECC. Many studies report children with multiple *S. mutans* strains (i.e., genotypes) having greater odds of developing ECC, studies investigating intraspecies interactions in dental caries are lacking. In this study, the impact of intraspecies interactions on cariogenic and fitness traits of clinical *S. mutans* isolates are investigated using *in-vitro* and *in-vivo* approaches.

Initially clinical *S. mutans* isolates of 10 children from a longitudinal epidemiological study were evaluated. *S. mutans* strains (G09 and G18, most prevalent) isolated from one child were used for subsequent analysis. Association analysis was used to determine if presence of multiple *S. mutans* genotypes within the first-year of colonization was associated with caries. Biofilm analysis was performed for single and mixed cultures to assess cariogenic traits, including biofilm biomass, intra-polysaccharide, pH, and glucan. Confocal Laser Scanning Microscopy (CLSM) and time-lapse imaging were used to evaluate spatial and temporal biofilm dynamics, respectively. A *Drosophila* model was used to assess colonization *in-vivo*.

Mean biofilm pH was significantly lower in co-cultured biofilms as compared with monoculture biofilms. Doubling of *S. mutans in-vitro* biofilms was observed by CLSM and *in-vivo* colonization in *Drosophila* for co-cultured *S. mutans*. Individual strains occupied specific domains in co-culture and G09 contributed most to increased co-culture biofilm thickness and colonization in *Drosophila*. Biofilm formation and acid production displayed distinct signatures in time-lapsed experiments.

**IMPORTANCE:** This study sheds light on the complex dynamics of a key contributor to early childhood dental caries (ECC) by exploring intraspecies interactions of different *S. mutans* strains and their impact on cariogenic traits. Utilizing clinical isolates from children with ECC, the research highlights significant differences in biofilm architecture and acid production in mixed versus single genotype cultures. The findings reveal that co-cultured *S. mutans* strains exhibit increased cell density and acidity, with individual strains occupying distinct domains. These insights, enhanced by use of time-lapsed Confocal Laser Scanning Microscopy and a Drosophila model, offer a deeper understanding of ECC pathogenesis and potential avenues for targeted interventions.

## INTRODUCTION

Dental caries is a complex, multi-factorial disease. Dental caries affects the oral health and overall health of over 80% of the human population worldwide.(1, 2) The Center for Disease Control and Prevention (CDC) (www.cdc.gov) lists dental caries as the most common chronic disease of children and adolescents. The World Health Organization reports that 60-80% of school children have dental caries in industrialized countries and predicts developing nations will see an increase due to consumption of fermentable carbohydrates and lack of proper fluoridation.(3) Severe early childhood caries (SECC) is of particular concern, because this form of the disease affects the health and well-being of children under six years of age due to extensive tooth decay.(4) SECC is highly prevalent in children from marginalized groups such as minorities, individuals of low socioeconomic status, and those with limited access to dental care.(4)

Many factors are thought to play a role in the disease initiation and progression, including the presence of specific oral bacteria. A primary focus in understanding the etiology of dental caries has been the search for microorganisms involved in the initiation, development and progression of the disease. Research suggests that oral disease may evolve from major homeostatic imbalances in the oral microbial community. Under certain conditions the ecology of the oral cavity selects for advantaged organisms that are conducive to a cariogenic state. Those dominate microorganisms include the mutans streptococci, lactobacilli, *Actinomyces*, bifidobacteria, and yeasts.(5, 6)

The primary bacteria typically associated with the initiation of early childhood dental caries are the mutans streptococci (MS), including *Streptococcus mutans* and *Streptococcus sobrinus.*(7–9) In the case of SECC, numerous studies have reported that the primary infectious agent is *S. mutans* and the presence of *S. mutans* is widely considered the most effective predictor of future caries incidence.(10) In our study population, children with higher *S. mutans* counts were 5.6 times more likely to develop caries and are significantly more likely to have higher caries scores.(11) The mutans streptococci are involved in the initiation of caries, creating an environment conducive to aciduric organisms colonization and demineralization of tooth surfaces.

Previously, we investigated *S. mutans* genetic diversity using different genotyping methods, repetitive extragenic palindromic-polymerase chain reaction (rep-PCR), multilocus sequence typing (MLST) and serotyping by PCR.(12–15) The scale and scope of this 8-year longitudinal epidemiological study affords a unique opportunity to determine the relationship between *S. mutans* genotypes and early childhood caries in a localized, high-caries risk population of African American children.(16) *S. mutans* strains are highly diverse in their phenotypic characteristics and cariogenic potential.(17, 18) Key pathogenic features include the ability to bind to teeth by formation of tenacious biofilms (i.e., dental plaque), to metabolize fermentable carbohydrates (especially sucrose) to produce organic acids, and to survive acidic environment (aciduricity).(17) Intracellular iodophilic polysaccharide (IPS) is an indicator of glycogen storage ability of *S. mutans* which allows for continued production of acid once carbohydrate sources are depleted.(19) Children in our study population with multiple *S. mutans* genotypes are 4.5 times more likely to have dental caries experience and younger children (age< 6 years) are 2.9 times more likely to have multiple genotypes.(20) Others have reported similar findings linking the presence of more *S. mutans* genotypes with higher caries scores.(21–27)

Despite literature supporting an association of intraspecies impact on caries potential, the majority of the research available has focused on evaluating either single or polymicrobial interspecies cultures or biofilms. Studies investigating the intraspecies interactions of multiple *S. mutans* genotypes and how these interactions contribute to caries virulence potential are lacking. In particular, studies investigating *S. mutans* intraspecies interactions in biofilms are crucial to examine how the presence of multiple *S. mutans* contribute to caries disease progression. Most studies investigating intraspecies interaction of *S. mutans* have largely focused on bacteriocins (i.e., mutacins in *S. mutans*) using stab agar antagonism assays and demonstrate that some *S. mutans* strains can have inhibitory effects on other *S. mutans* (28–30). This poses the question: is the presence of multiple *S. mutans* mutualistic or antagonistic in forming the cariogenic environment? Based on our previous association data for our study population, we hypothesized that the presence of multiple *S. mutans* genotypes contribute significantly to cariogenic traits and may provide a mutualistic benefit for *S. mutans* fitness. In this study, we investigate the impact of multiple *S. mutans* genotypes on cariogenic potential using *in-vitro* (biofilm) and *in-vivo* (*Drosophila* model) approaches.

## RESULTS

### Presence of multiple *S. mutans* genotypes within the first year of detection is significantly associated with caries

Based on the summary of longitudinal data (8-years), we previously reported that younger children within our study population were more likely to have multiple genotypes and that there was a significant association between having multiple genotypes and dental caries(20). Here, we determine if having multiple *S. mutans* genotypes at initial detection of *S. mutans* or within the first-year of detection is associated with dental caries (decayed, missing, filled, surfaces-DMFS score>0 within 6-years), a cross-sectional analysis was performed. Children with multiple genotypes at initial detection were not significantly associated with developing caries. This is logical as there is a time lapse from initial colonization to development of detectable caries.

However, the presence of multiple genotypes within the first-year of detection was found to be significantly associated with developing ECC (**Table 1**) highlighting that early colonization with multiple *S. mutans* is a risk factor for caries. For children with multiple genotypes, the mean number of genotypes was 2.7 (range 2-6 genotypes) within the first-year of detection. Of the 14 caries-free children in both groups (10 single or four multiple), 12 (86%) children did not maintain persistent colonization of *S. mutans* for the duration of the study (6-years).

**Table 1.**
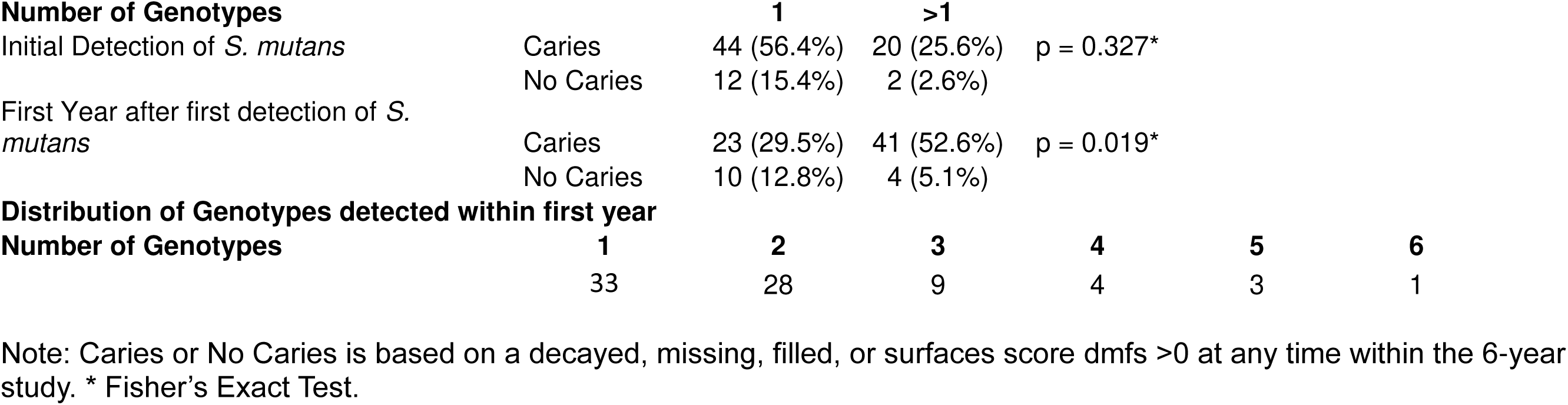
Presence of multiple *Streptococcus mutans* vs caries (n=78)

### Biofilm pH is significantly lower in co-cultured *S. mutans* biofilms

To investigate the effects of co-culture of different genotypes of *S. mutans* from the same individuals, we evaluated static biofilms for key cariogenic traits (biomass, pH and IPS) for 10 children with either two or four *S. mutans* genotypes (termed genotype “profiles” for each child). Co-cultured isolates from 9/10 children showed a trend that co-culture biofilms have lower biofilm pH compared to mean mono-culture biofilms as indicated by increased fluorescence of the pHrodo probe (**Fig. 1A**). Profile-5 (2-genotypes) and Profiles-1, 7, 8, 9 (4-genotypes) had significantly lower mean biofilm pH for the co-culture biofilms than combined mono-cultures. Overall mean biofilm pH for the population (10 children) of the co-cultured showed a significant increase in pHrodo fluorescence, indicating lower biofilm pH and significant greater cell density by increased Syto9 fluorescence (**Fig. 1B**). No overall significant differences in biofilm biomass or IPS was observed for the population (**Fig. S1**). For Profile-5 (subject C-232), selected for subsequent analysis, the mean co-culture biofilm consistently had significantly lower biofilm pH than mono-culture (**Fig. 1C**).

**FIG 1.**
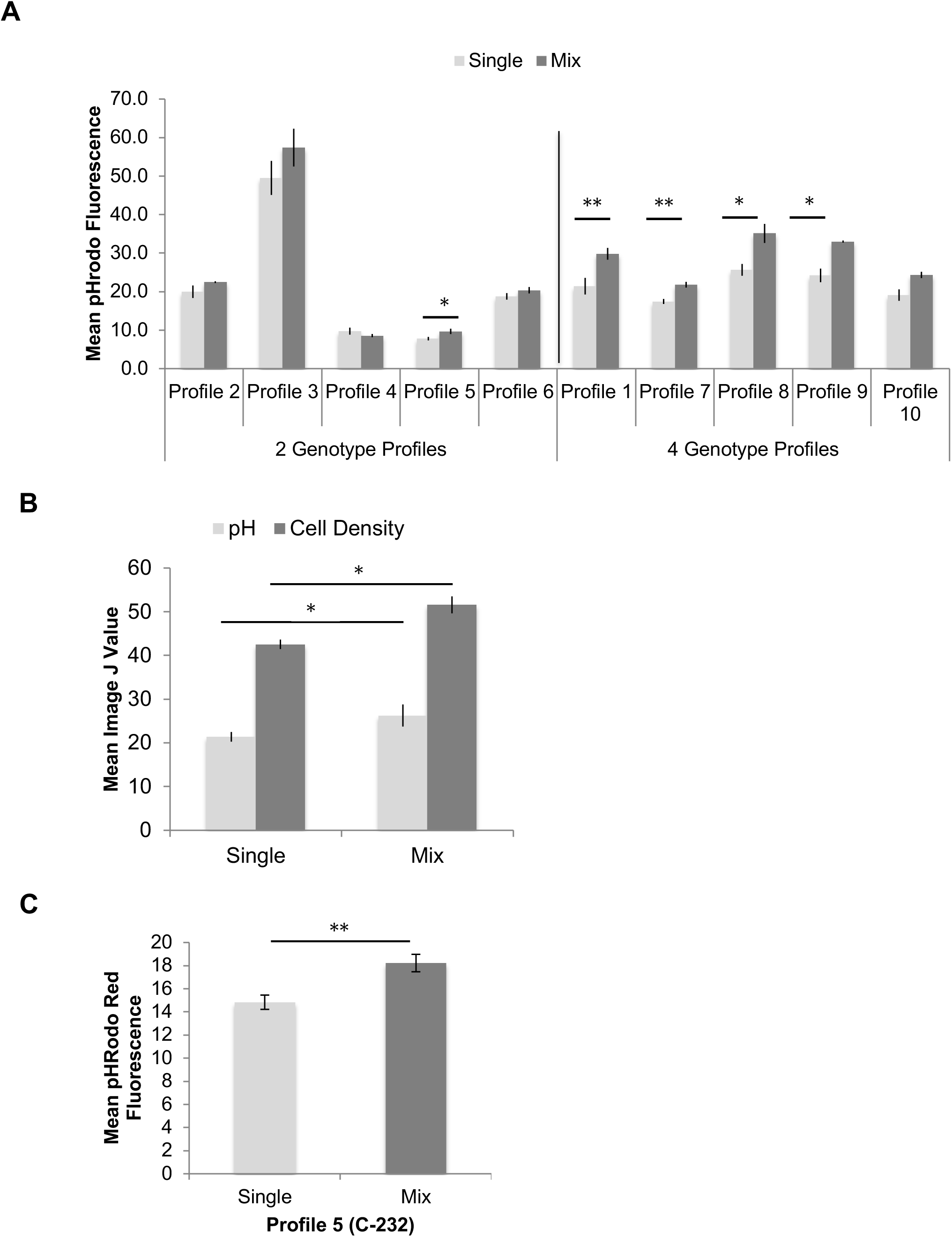
Impact of mono-cultured (Single) vs co-cultured (Mix) *S. mutans* on biofilm pH. (A) Children with mixed genotypes demonstrate a trend of decreased mean biofilm pH in mixed versus single strain cultures from 9 out of 10 children. Lower pH is shown by increased fluorescence of pHrodo dextran conjugated probe. Mean fluorescence data obtained using Image J. Single experiment 3 technical replicates to illustrate population trend. **<**Profile= is *S. mutans* genotype profile for each child. (B) Overall mean biofilm for all 10 children shows significantly lower biofilm pH and higher cell density by mean Image J analysis of fluorescence. (C) *S. mutans* from child 232 consistently demonstrates significantly lower biofilm pH for the mixed strains than single strains alone. Data from at least 3 independent experiments with 3 technical replicates each. Isolates from Profile 5 (C-232) was selected for continued study because it presented with G09 and G18, the most common genotypes observed in this high caries risk population. Standard error bars shown. * P<0.05, ** P<0.01, ***P<0.001

Profile-5 consisted of *S. mutans* genotypes G09 and G18, previously identified by rep-PCR (**Fig. 2)**. No significant difference was observed in biomass or IPS between mono- and co-cultured strains, although the mix did trend higher for IPS. Biomass was significantly lower for both mono-cultured and co-cultured biofilms when compared to *S. mutans* UA159 control, which is consistent with previously reported data for the representative library G18 strain (UAB-10) (**Fig. 2A**).(16) However, IPS was significantly higher for mono- and co-cultured biofilms of clinical isolates as compared to *S. mutans* UA159 (**Fig. 2B**) indicating greater glycogen storage in the clinical isolates.

**FIG 2.**
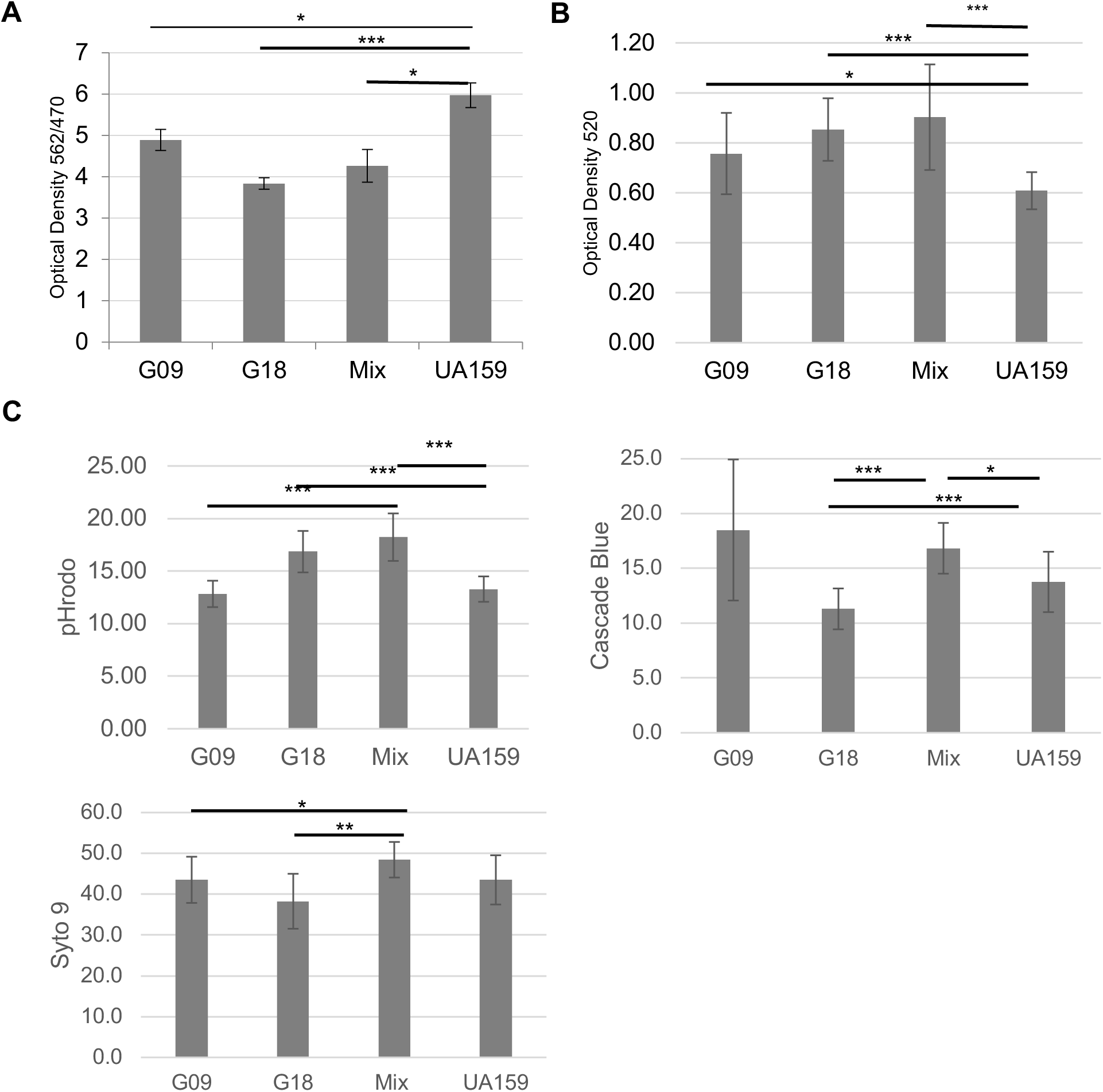
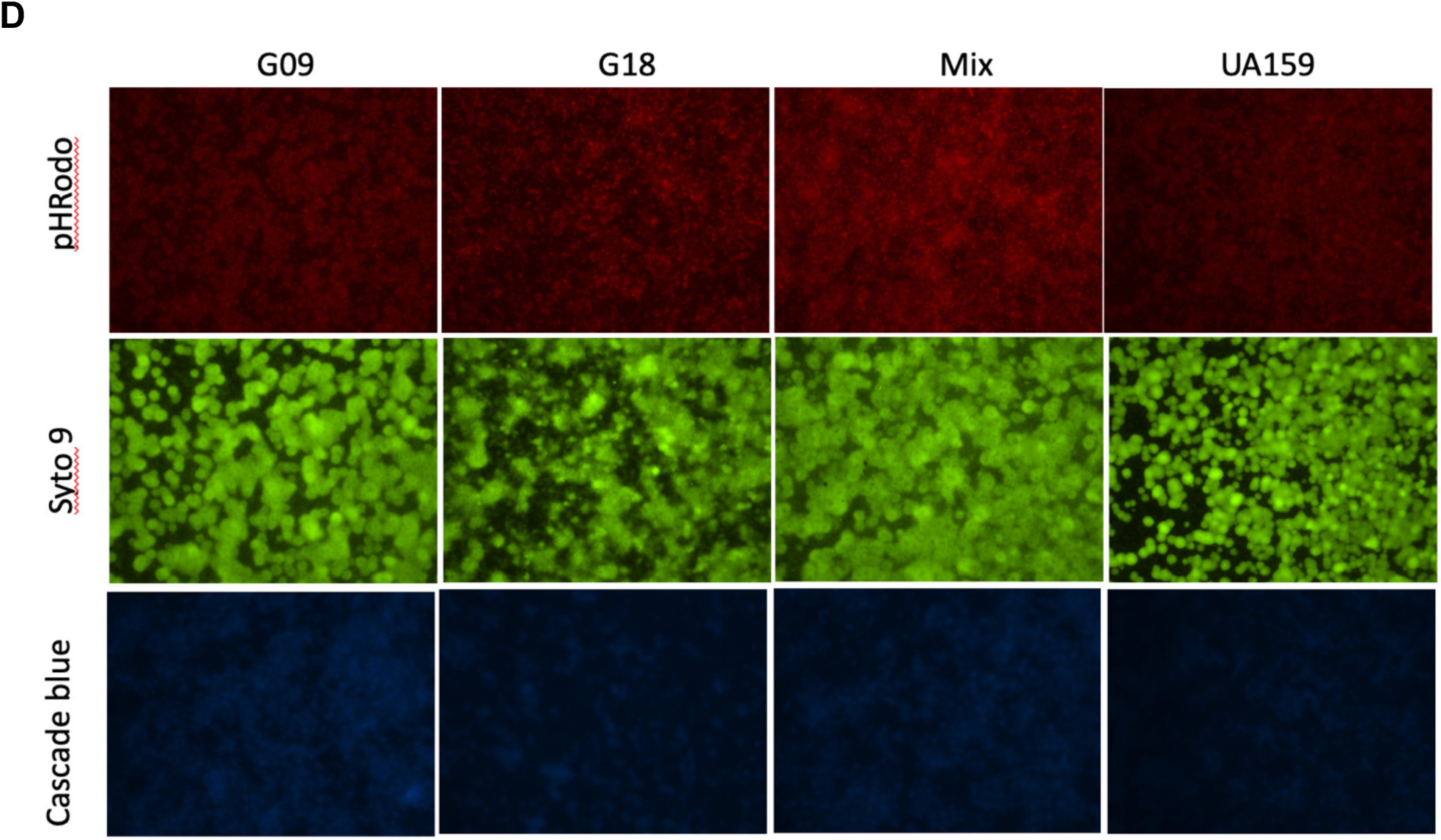
Effect of co-cultured *S. mutans* (Mix, G09 and G18) versus mono-cultures (G09 or G18) on virulence properties in-vitro. Both G09 and G18 were from child C-232. (A) Mean biofilm mass by crystal violet assay normalized to cell growth (OD_470_) indicates no significant difference between mono- and co-cultures biofilms. (B) Mean intracellular polysaccharide by iodine assay demonstrates no significant difference between mono- and co-culture biofilms. (C) Mean Image J values for biofilm pH (pHRodo, red), glucan (cascade blue), and cell density (Syto9, green) reveal increase in pHrodo fluorescence and cell dentisty between mono- and co-culture biofilms. (D) Representative images of biofilm pH (pHrodo red), glucan (cascade blue) and cell density (Syto9 green) shown with 10x magnification. Increase brightness of the mix (co-cultured) biofilm is markedly brighter. All biofilms were performed in Todd Hewitt Broth + 1% sucrose. N=3 with a minimum of three technical replicates. Standard deviation error bars are shown. * P<0.05, ** P<0.01, ***P<0.001 by Student9s *t* test.

For biofilm pH of Profile-5, co-cultured biofilm was more acidic (indicated by increased fluorescence of pHrodo) than both mono-culture biofilms (**Fig. 2C and D**). Both G18 mono-culture and co-cultured biofilm were significantly more acidic than *S. mutans* UA159 which is consistent with previously published findings for another G18 strain, UAB-10.(16) When the combined mean of the mono-cultures are compared to the mean co-cultured, the mix remains significantly more acidic (**Fig. 1C**). Cascade blue is a dextran conjugated dye used to label extracellular glucans (**Fig. 2C and D**). Glucan was lower for G18 was significantly lower when compared with the co-cultured biofilms. This reduction in glucans for G18 did not impact the pHrodo data for G18, indicating the acidic phenotype observed in co-culture was not due to variable glucan production between strains. Cell density (Syto9) in 2D imaging was consistently significantly higher in the co-culture biofilm as compared to mono-cultured biofilms indicating greater cell growth within the mix.

### Co-culturing *S. mutans* significantly increases *S. mutans* cell density and biofilm thickness by CLSM

To investigate the impact of co-culture on the biofilm thickness and architecture, *S. mutans* G09 and G18 were genetically modified with mCherry red and green fluorescent protein (GFP), respectively, for CLSM analysis. The mean fluorescence of mono- and co-cultured biofilms were comparable for individual strains while G18 was significantly lower than UA159 (**Fig. 3A**). However, considering biofilms were inoculated with comparable concentrations of *S. mutans* (mix composed of 50% each strain), there was a notable doubling effect observed in the co-cultured total fluorescence suggesting increased cell density. This was also observed when measuring biofilm thickness, however G09 showed significantly greater biofilm thickness in the mix than its mono-culture (**Fig. 3B**). In comparison with UA159, both mono-cultures had significant less thickness than UA159. The statistically significant doubling of *S. mutans* in the co-cultured versus mono-cultures and UA159 is best illustrated with mean volume/area as this represents the total area of the fluorescence of *S. mutans* within the biofilm (**Fig. 3C)**.

**FIG 3.**
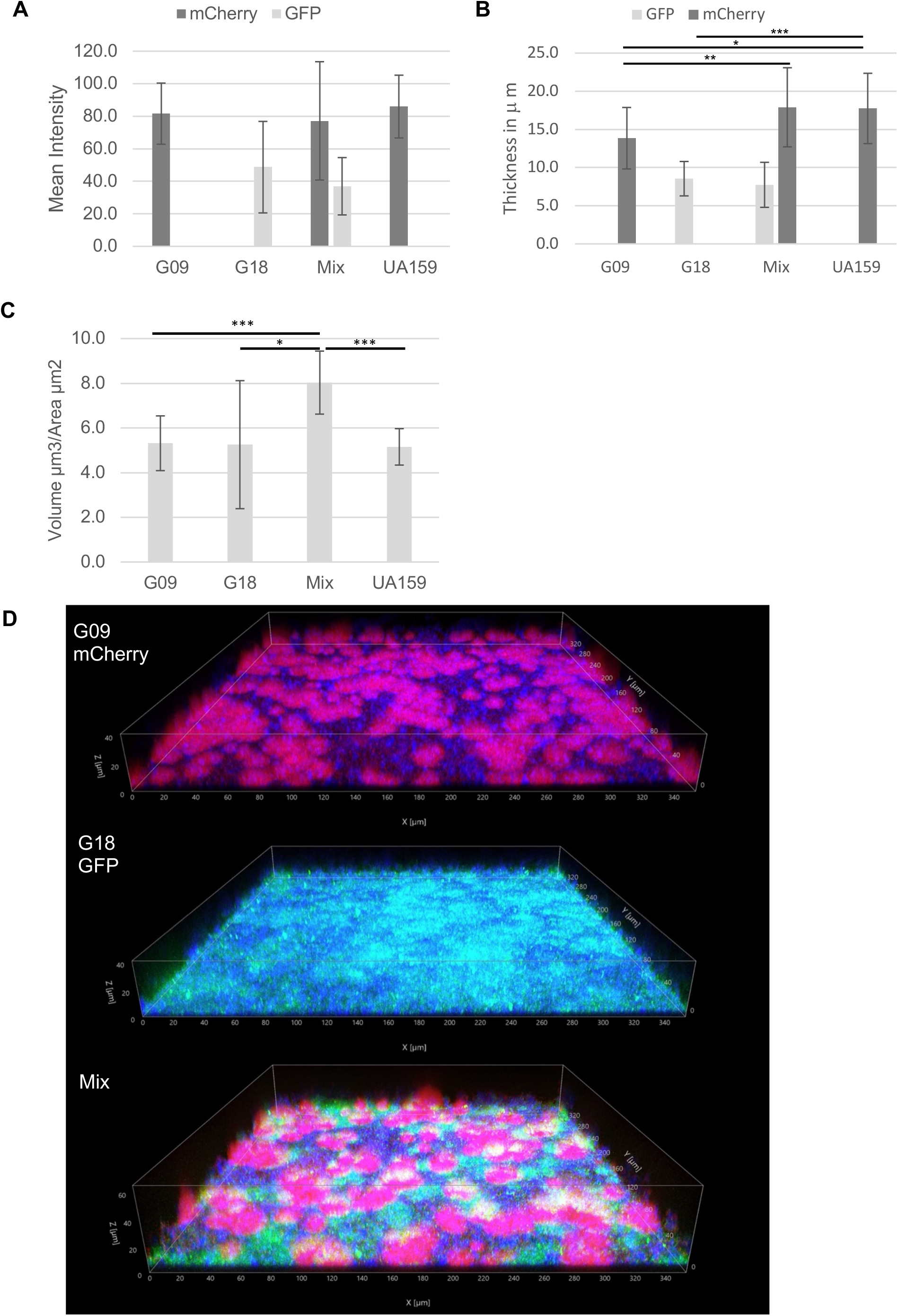
CLSM imaging and quantification of *S. mutans* single and mixed biofilms. (A) Mean fluorescence intensity for *S. mutans* by CLSM. (B) Mean *S. mutans* biofilm thickness by CLSM. Both fluorescence intensity and thickness show a doubling of total *S. mutans* within the mix, though only G09 in the mix shows a significant difference in biofilm thickness. (C) Average volume/area demonstrates significant doubling effect of *S. mutans* in mix versus single culture biofilms. (D) CLSM image for single G09 (mCherry red), G18 (GFP green), and mix (red & green) illustrating shift in spatial architecture and biofilm heights of *S. mutans* in single vs. mixed biofilms. Cascade blue was used to image extracellular glucan matrix. N=3 with 5-7 technical replicates per experiment. Note: Video image of single and mixed biofilm CLSM is available in supplemental materials online. * p <0.05, ** p<0.01 and *** p<0.001

The shift in spatial arrangement and biofilm thickness in mixed biofilm vs mono-culture biofilms is illustrated in **Figure 3D and Movie S1**. In mono-culture, G09 (red) forms large aggregates with noticeable glucan matrix (blue) forming the extracellular space between while G18 demonstrates a confluent lawn phenotype with minimal visible glucans. However, in co-culture G09 forms larger and taller aggregates (**Fig. 3D and Fig. S2**), which collapse inward on themselves, forming volcano-like structures with negative space in the underlying glucan layer (**Movie S1**). Interestingly, each strain occupies a specific domain with no apparent overlay.

It is noteworthy that mCherry was not responsible for the aggregate phenotype observed in G09. A subsequent experiment of G09 with GFP displayed the same phenotype (not shown). However, labeling G18 with mCherry did promote an aggregate phenotype (data not shown) indicating that mCherry can alter biofilm formation in some clinical strains of *S. mutans*. Thus, the use of an alternate fluorescent protein is encouraged as a control when possible.

By CLSM analysis, glucan estimated by cascade blue fluorescence was significantly greater for G18 on average as compared to G09 with G18 and co-cultured biofilms produced significantly more glucan than UA159 (**Fig. S3A**). No significant difference was observed in glucan biofilm thickness between mono-culture, mix and UA159 (**Fig. S3B**). The fluorescent of glucan by CLSM seemingly contradicts the 2D imaging data (**Fig. S3C**). However, 3D imaging used more sampling sites and multiple z-stack layers making it is a more reliable measure to report the *in-situ* glucans.

Given difference in pH, cell density, and thickness, it was crucial to confirm that the inoculum of *S. mutans* used were comparable between mono- and co-cultures at set-up. To evaluate this, G09 and G18 were genetically modified with different antibiotic resistance genes and differential plating was used at biofilm set-up and after biofilm formation. No significant difference in the total number of *S. mutans* used to inoculate the biofilms were observed using colony forming units (CFUs/mL) plating on antibiotic selective media for either pre-plate or post-plate (**Fig. S4**). The distribution of both strains within the co-culture biofilms both pre- and post-were comparable with each comprising approximately half the total *S. mutans* in the mix.

### Different *S. mutans* clinical strains display distinguishable biofilms and acid formation phenotypes

Since co-cultured biofilms displayed a dramatic difference in biofilm structure, we next evaluated the formation of the biofilms using time lapsed imaging for 24-hours. Using this approach, it was possible to observe biofilm formation dynamics for G09 and G18 as early as 3-4 hours (**Fig. S5**). Interestingly, biofilm formation was clearly distinguishable between the two *S. mutans* strains in single and co-cultured mix biofilms over the first 12-hours. This difference was best observed in the contrasted oblique illumination mode (**Fig. S6 or Movie S2**), G09 displays dense microcolony-like mini-aggregates early in biofilm formation while G18 forms thinner, broader “fish net” networks of streptococccal chains. These distinct architectures are also observed in the fluorescence mode as the biofilm develops (**Fig. S6 and Movie S2**). As previously observed in the CLSM, the two strains occupy distinct domains within the biofilm with no appreciable overlap observed.

Since biofilm formation was distinguishable between G09 and G18, we subsequently evaluated biofilm pH dynamics with pHrodo red using 30-hour time lapse imaging with both strains labeled with GFP. Time was extended to 30-hours to better capture the second pH fluorescence increase. The pHrodo fluorescent intensity continually increased over time with two distinct phases at 10-11 hours and 24-26 hours (**Fig. S7A**). GFP fluorescence peaked between 8-9 hours then decreased for several hours until GFP intensity partially recovered and plateaued around 25-26 hours (**Fig. S7B**). The decrease in GFP fluorescence may be due to GFP’s known sensitivity to low pH or *S. mutans* acid tolerance, but the overall biofilm adapts to the new aciduric environment.

Fluorescent intensity of pHrodo consistently peaked within the 2-hours following the GFP fluorescence peaks.

### Co-culture of *S. mutans* improves *S. mutans* colonization *in-vivo*

A fly model was used to evaluate *in-vivo* colonization of single or co-cultured *S. mutans*. Using the capillary feeding *Drosophila* model illustrates how co-cultures of *S. mutans* influence colonization within the *Drosophila* as compared to mono-cultures. The consumption of bacteria in 5% sucrose was comparable with no significant difference when comparing mean feedings for either day or night feeding across strains (**Fig. 4A**). There was a significant increase in day consumption for both single strains as compared to night feeding. *S. mutans* G09 colonization within *Drosophila* was consistently greater when G18 was also present (**Fig. 4B**). This is consistent with what observations by CLSM. Interestingly, when the co-culture was plated on THA without antibiotics (total *S. mutans* count), the CFUs were equal to the mean of the mono-cultures. However, growth on the antibiotic selective media were more consistent with CLSM, exhibiting the doubling of total *S. mutans*, suggesting the use of antibiotic specific media is required to improve recovery and obtain accurate differential counts.

**FIG 4.**
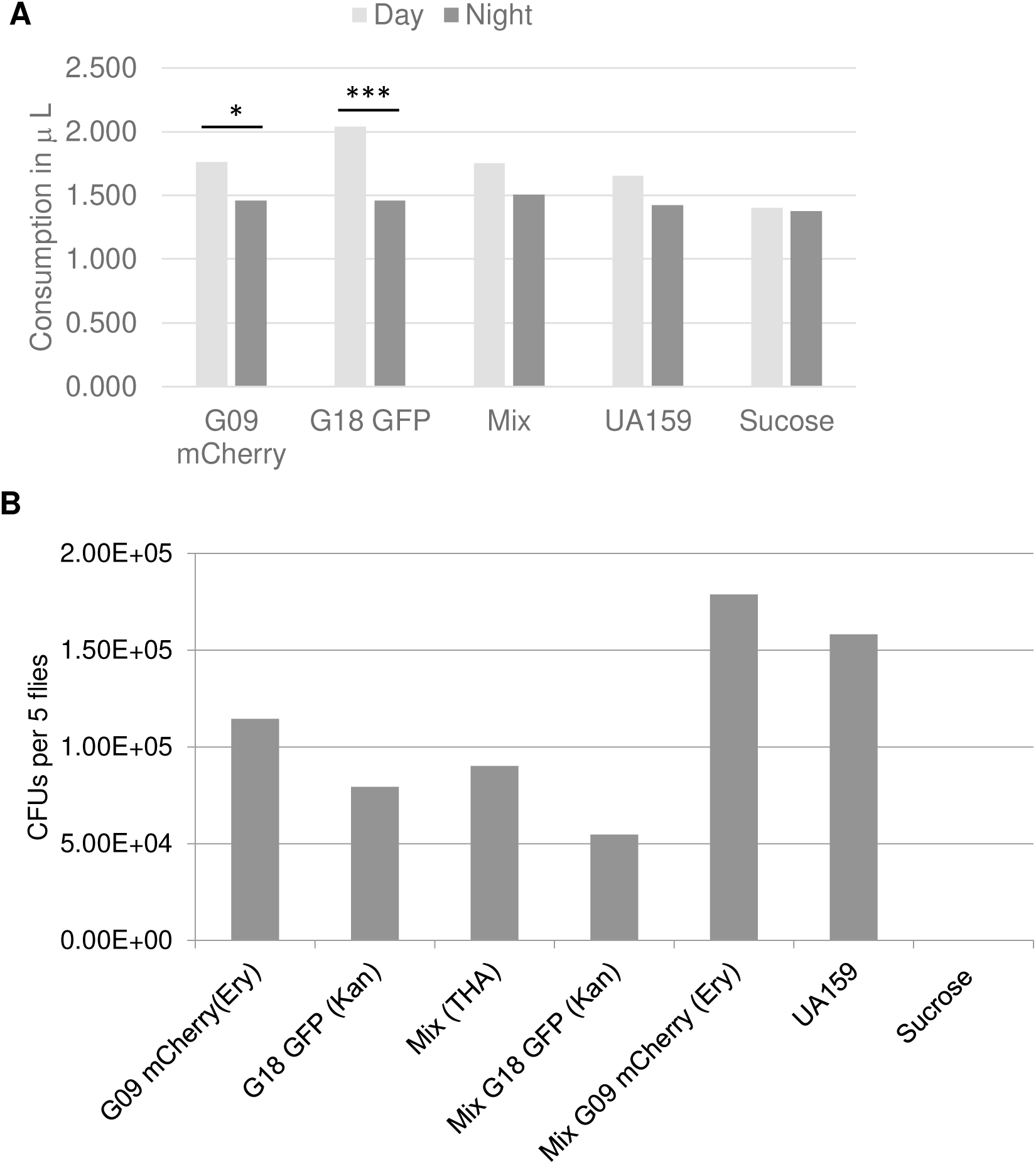
Mean *S. mutans* consumption and colonization in *Drosophila* with mono- and co-cultures. GFP = green fluorescent protein, mCherry = red fluorescent protein. (A) Normalized mean consumption of bacterial suspensions in 5% sucrose for 7 flies over 3 days. Day = 8 hours, Night = 16 hours/2. No significant difference was observed for the mean consumption over 3 days. UA159 and sucrose are positive and negative controls, respectively. (B) Mean colony forming units (CFUs) per 5 flies differentially plated on Todd Hewitt Agar with and without antibiotics shows increased colonization of G09 with decreased G18 in the co-cultured as compared with mono-cultures. N=3 with 2 technical replicates each. Error bars are not show as variance is high between experiments, however trend remains consistent. * P<0.05, ** P<0.01, ***P<0.001 by Student’s *t* test.

## DISCUSSION

Numerous associations studies report the presence of multiple *S. mutans* strains is associated with greater childhood caries, (20–27) but studies exploring intraspecies interactions between *S. mutans* are limited. While interspecies studies are important to the study of dental caries, it is equally important to understand how intraspecies can also contribute. The oral cavity is a diverse ecological environment that is constantly under a variety of environmental stresses. (31) Therefore, oral bacteria, especially *S. mutans* are particularly adept at adaptive horizontal gene transfer making for rich accessory genomes.(16, 32–35) In this study molecular biology approaches were applied to determine if the presence of multiple *S. mutans* are directly related to changes in measurable caries virulence traits.

In the initial population study for 10 children, it was notable that more of co-cultured biofilms for 4-genotype children were significantly more acidic as compared to the children with 2-genotypes (Fig. 1A). This population-based initial approach indicates that the presence of more *S. mutans* can contribute to an acid environment.

Despite having less biofilm biomass than the known cariogenic *S. mutans* UA159 strain, the biofilm analysis for C-232 isolates suggests that G18 contributes most to the significantly acidic biofilm pH (**Fig. 2C and D**) while G09 significantly contributes more to biofilm thickness (**Fig. 3B**). When co-cultured these two stains provide a mutually beneficial advantage in overall colonization, particularly for G09, leading to greater cariogenic potential over mono-cultures. Since both pHrodo and cascade blue are dextran conjugated, and glucan was reportedly lower in G18 mono-culture, the actual increase in biofilm pH may be underestimated compared to G09 and mix.(36) This finding is different than we previously reported for the representative library strain G18 (UAB-10) which produced significantly more glucan than UA159 control.(16) Conversely, the glucan data from the CLSM fluorescence data showed glucan for G18 trended higher than the mix (**Fig. S3A**). However, data from the CLSM is a more reliable imaging *in-situ* as noted in the results.

The increase in cell density of *S. mutans* observed in the 2D-biofilm imaging with Syto9 (**Figs. 1B and 2C**) together with the doubling effect in both CLSM analysis (**Fig. 3**) and drosophila model (**Fig. 4**) for the co-cultured samples supports that the presence of multiple *S. mutans* is mutually beneficial to the formation of biofilms. Even though there was no significant difference in biofilm biomass (**Fig. 2A**), the increase in biofilm thickness and volume by CLSM (**Fig. 3B and C**) demonstrates the formation of more robust biofilms can improve colonization and persistence. Although, differential plate enumeration of CFUs/mL did not show a significant difference for post-biofilms analysis (**Fig. S4**), several factors may account for this difference, most notably the challenge of disrupting large biofilm aggregates efficiently.

Some studies have evaluated the spatial structure of oral biofilms using *in-vitro* and *in-vivo* polymicrobial models, demonstrating that oral biofilms form highly organized, spatially structured communities.(37–41) The present study differs in its CLSM analysis of *S. mutans* intraspecies impact on spatial distribution and arrangement. Early colonization of *S. mutans* remains one of the best clinical indicators of ECC, especially in our study population.(11) Given the high number of children within our study population with multiple *S. mutans* genotypes, it was worth investigating how *S. mutans* intraspecies interactions shape the early development of oral biofilm and the spatial landscape. The CLSM analysis supports that co-cultured *S. mutans* biofilms leads to increased biofilm thickness (**Figs. 3B, 3D, S2 and Movie S1**). Furthermore, CLSM and time lapsed imaging revealed that clinical *S. mutans* G09 and G18 each demonstrated unique phenotypes, occupied specific domains within the biofilm, and have no observable colocalization (**Fig. S5, Movie S2**). The domed phenotype previously reported by Kim and Koo (38) was more similar to what was observed for the G09 phenotype in this study and differs significantly from the lawn phenotype observed for the G18 strain. Greater biofilm height from *S. mutans* G09 coupled with the fuller coverage of G18 contribute significantly to the surface area available for other organisms to attach, increasing overall cariogenic potential of co-cultured *S. mutans* biofilms. Future studies are planned to evaluate how intraspecies *S. mutans* interactions impact the spatial landscape when other species are present.

Although *S. mutans* UA159 has been widely studied and is known to be cariogenic in a rat model, it is crucial to understand that other clinical *S. mutans* can contribute to caries in a very different manner than UA159. It has been well documented that clinical *S. mutans* can differ considerably in their cariogenic phenotypes.(17, 32, 42) It is important to note biofilm experiments should be tested with other clinical strains, including those showing inhibition against other *S. mutans,* to provide a more comprehensive understanding of intraspecies interactions among clinical isolates.

The time-lapsed data included in this study also provides a valuable new information on how acid is formed (**Fig. S7**). The observed bi-phasic effect in the pHrodo suggest *S. mutans* may be going through a period of equilibration in response to the initial acid production, possibly through the activation of genes related acid tolerance response (ATR) due to acid stress.(17, 43, 44) Future studies needed to elucidate this phenomenon.

The finding that the presence of multiple *S. mutans* within the first-year of detection is significantly associated with having caries is informative, indicating the presence of multiple *S. mutans* is linked and likely contributes to early onset of ECC within the first-year of colonization. Among caries free children, the majority (86%) did not show stable colonization of *S. mutans*. That is, *S. mutans* strains were acquired and lost; or colonized in subsequent periods with a different strain which were later lost. This finding is particularly interesting in that these children would make excellent subjects for subsequent host immunology and microbiome studies to determine why *S. mutans* was unsuccessful in colonizing the oral cavity of these children.

In summary, when multiple *S. mutans* strains G09 and G18 are present, there are significant increases in cariogenic properties including colonization, surface area coverage, cell density, and acid production. Given the high prevalence of these strain types within our high-caries risk population and the large percentage of children with multiple strain types within the first-year of detection of *S. mutans*, suggests having multiple genotypes, particularly *S. mutans* G09 and G18 are strong risk factors for early childhood caries.

## MATERIALS AND METHODS

### Bacterial Strains

*S. mutans* isolates from African-American children from a previous 8-year longitudinal study in a high-risk caries population were used. Since longitudinal data of acquisition of *S. mutans* genotypes was available, this provides a key advantage in the present study since a patient-based model was used (i.e., actual strains from specific caries-active children were used).

Previously, we reported an overview analysis of single vs. multiple genotypes within this population with details on collection methods and rep-PCR analysis.(20, 45) For the present study, we provide additional analysis on the relationship of having multiple *S. mutans* at initial detection of *S. mutans* or within the first-year after initial detection with caries (yes/no) for the younger cohort of children (Cohort 2) who had at least 7 isolates (n=78) analyzed by rep-PCR. Caries was defined for a child with a decayed, missing, filled surfaces (dmfs)>0 at any time during the previous study (6-years). (46)

Preliminary analysis of biofilms was performed with *S. mutans* isolates from 10 caries-active children (5 children presenting 2-genotypes, 5 children presenting 4-genotypes during the previous study) (**Table 2**). One child (C-232, Profile-5) with 2-genotypes was selected for further study. This child presented with *S. mutans* genotypes G18 and G09, the two most prevalent genotypes observed in the larger epidemiological study of this high-caries population.(20) *S. mutans* UA159 was used as a positive control on all experiments.

**Table 2.**
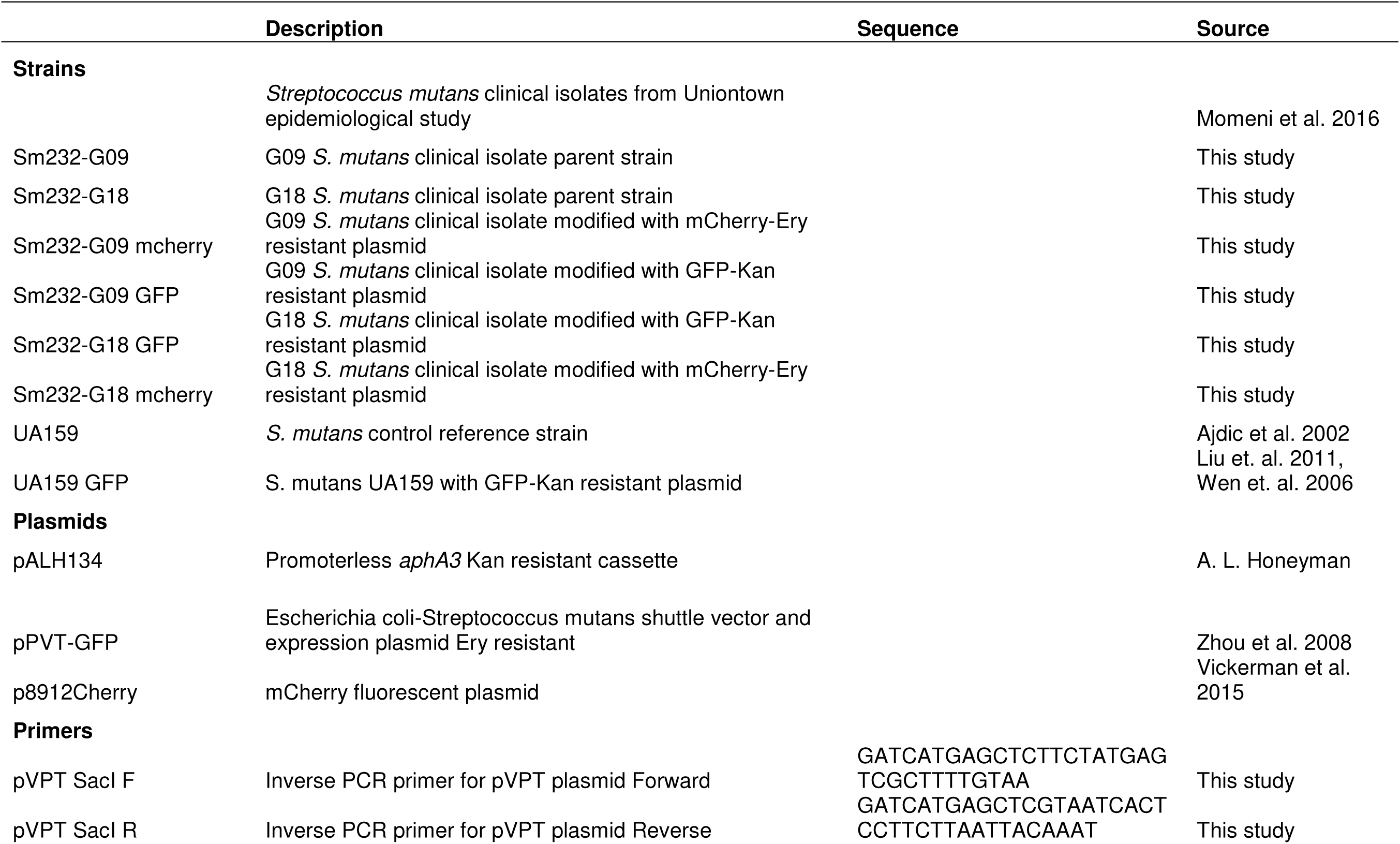
List of bacterial strains, plasmids, and primers used in this study.

### Mutant Constructions

To determine strain prevalence within the mixed biofilm matrix and biofilm spatial arrangement by confocal microscopy the *S. mutans* G18 and G09 strains from C-232 were genetically modified with plasmids carrying different fluorescent proteins and antibiotic resistance genes (green fluorescent protein pVPT +Kan^R^ and red fluorescent protein mCherry+Ery^R^, respectively) (**Table 2**).(47–49) To make the pVPT-GFP+Kan^R^ plasmid, the erythromycin gene of the pVPT+GFP plasmid was removed using reverse PCR with the pVPT SacI forward and reverse primers (**Table 2**). Next, PCR ligation mutagenesis was used to insert a nonpolar kanamycin resistance cassette obtained by enzyme digestion from pALH124 using SacI restriction enzyme. Ligation product was transformed into *E. coli* cells and kanamycin resistant transformants were PCR screened to confirm mutation. Plasmids were confirmed using sequencing prior to transformation into *S. mutans* strains. Antibiotics were used in the following concentrations: 1mg ml^-1^ kanamycin and 10μg ml^-1^ erythromycin for *S. mutans* and 50μg ml^-1^ kanamycin for *E. coli*.

### Biofilm Analysis

*In-vitro* static biofilm analysis was performed as previously described.(16) Briefly, fresh from frozen stocks were plated on Todd Hewitt Agar (THA) and grown anaerobically (10% hydrogen, 10% carbon dioxide, 80% nitrogen) for 48-hours at 37°C then sub-cultured in Todd Hewitt Broth (THB) to optical density OD_600_≍0.5 before diluting 1:1,000 in THB+1% sucrose. Samples were prepared for individual mono-cultures and co-culture (mix of two or four *S. mutans* together). For single-cultures, 5μl of subculture was added to 5ml THB+1% sucrose for biofilm set-up. For co-cultured mixed samples, 10μl of each sub-cultured strain was mixed, then 5 μl of this mixture was used for biofilm set-up, such that all biofilms were inoculated with comparable volumes of *S. mutans*. For the pH assay, pHrodo red or cascade blue dextran conjugated probes (Molecular Probes, Invitrogen, Carlsbad, CA) were added 1:1,000 prior to biofilm set-up to monitor pH and glucan production, respectively. A 96-well flat-bottom microplate was used with 200μl per well. All Biofilm plates were incubated for 16-hours in 5%CO_2_ incubator at 37°C under static conditions. Only one biological replicate was performed in the initial analysis for the 10 children (triple technical replicates), as the purpose was to demonstrate a trend within the overall population. For all subsequent analysis of C-232 (Profile-5), a minimum of 3 biological replicates, 3 technical replicates each were performed.

Biofilms analysis was performed initially to assess biomass(crystal violet assay), intracellular iodophilic polysaccharide(IPS, to assess glycogen by iodine assay), and pH(pHrodo/cascade blue/Syto9) for the 10 children as described.(16, 19, 50–53) OD*_562_*and OD_520_ were measured for biomass and IPS, respectively. To determine biofilm pH, wells were manually surveyed and 2D representative images collected at 10x using a fluorescent microscope (Nikon TE2000-E inverted scope). ImageJ was used to calculate mean values of the fluorescence to determine pH(pHrodo), glucan(cascade blue) and cell density(Syto9).

### Confocal Laser Scanning Microscopy (CLSM) Analysis

Biofilms were prepared as described above, except in eight-well ibidi treated slides (ibidi, Fitchburg, WI) with 300μl per well in duplicate wells. Overnight biofilms were washed 3x with 1x phosphate buffered saline (PBS). All confocal was performed with 3 biological replicates, with a minimum of 5 image sites collected per sample.

Biofilms were imaged on a ZEISS LSM 880 or LSM 980 laser-scanning confocal through a 40x1.2 LD LCI Plan-Apochromat water immersion lens and excitation and emission parameters suggested by the manufacturer for mCherry, EGFP, and Cascade Blue. Image analysis of average intensity for all three channels in volume segmentations was performed in Bitplane Imaris v10.0.1. The average height of the biofilms was assessed from orthogonal projections spanning the full depth of the field of view.

### Time-Lapsed Biofilms Analysis

To examine the biofilm architecture formation over time, biofilms were prepared as described above for CLSM and imaged every hour for 24-hours using a Zeiss CellDiscoverer7 (CD7) instrument using a 50x PlanApo 1.2NA water immersion lens in combination with a 0.5 tube lens and detection on a Zeiss Axiocam 506m camera at a pixel resolution of 0.181μm per pixel and a field-of-view of 320μm by 285μm. Imaging environment was kept at a constant 37°C and 5%CO_2_. A 40μm z-stack at 1μm interval was acquired every hour for 24-hours with two fluorescence channels (mCherry and GFP) as needed, as well as a brightfield image contrasted through oblique illumination. The fluorescence and brightfield channels were processed separately for each position over time. Background in fluorescence channels was equalized and subtracted using a rolling ball algorithm, and channel crosstalk was reduced using linear unmixing. The bottommost 6 slices of the fluorescence channels were orthogonally projected for maximum intensity in each pixel and overlaid with the most contrasted transmitted image from the z-stack prior to export into a movie file.

To investigate the formation of biofilm pH over time, *S. mutans* G09 GFP and G18 GFP were used with pHrodo red dextran conjugated probe to determine pH with the methods described above with imaging performed every hour for 30-hours. All 24-30 hours experiments were performed in duplicate on two independent experiments with 3-4 sites per well and cascade blue was not used in these experiments.

### Differential Plating

To determine the distribution of G18 and G09 within the single and mixed biofilms differential plating on specific antibiotic agars was used. “Pre-plate” was from the original culture used to inoculate the biofilms to determine starting amounts of *S. mutans* were accurate and comparable between single and mixed cultures. “Post-Plate” was from 16-hour biofilm, washed three times with sterile 1x PBS, manually dislodged, and briefly sonicated. All samples were serially diluted, then 100 μl plated in duplicate on THA with respective antibiotics. Colony forming units (CFUs/mL) were manually enumerated.

### *Drosophila* Colonization Model

A *Drosophila melanogaster* model was used to determine colonization *in-vivo* for the single and co-cultured *S. mutans*. This model has been commonly used to determine colonization of *S. mutans* and other bacteria.(47, 53–57) Flies were orally infected by modified capillary feeding system (**Fig. S1**).(58) Seven male Canton-S flies (1-3 day old) were treated with antibiotics (Jazz Mix Drosophila food with erythromycin, vancomycin, and ampicillin 50μg/mL) for 2-days. Fresh overnight (first feeding) and mid-log sub-cultured *S. mutans* (OD_600_ ≍ 0.5, second feeding, 8-hours after first) were used. Respective antibiotics were used to maintain plasmids in both overnight and subcultures. Aliquots of 1mL for single culture and 500μL for of each culture for mixed sample (total volume also 1mL) were centrifuged at 10,000 rpm for 3-min and bacterial pellets washed 2x with 1x sterile PBS to remove residual media and antibiotics.

Harvested cells were resuspended in 100μL 5% sucrose and 0.5μL of suspensions were added to calibrated micro-capillary tubes (Miles Scientific, Newark, DE) inserted via pipette tips through cotton vial caps. Flies were incubated at room temperature and protected from direct light. Flies were fed by capillary feeding for 3-days then knocked out in -20 freezer for 1-hour. Next, five flies were briefly sterilized with 70% ethanol and washed 3x with sterile PBS. Flies were pulverize in 200μL sterile PBS. Serial dilutions of homogenate were spread on respective THA with antibiotics. Plates were incubated 48-hours at 37°C in 5% CO_2_ incubator and CFUs enumerated.

Some challenges to the *Drosophila* capillary feeding model were specific to this study. The single feeding model did not work well with *S. mutans* since these readily form biofilms within the capillaries over time which blocked flies from access to the food (bacteria in sucrose), so a twice daily feeding was performed with the second, longer feeding (16-hours) being made with fresh sub-cultured *S. mutans* cells. This appeared to minimize biofilm formation within the capillaries and allowed continued access to the bacterial/sucrose source. An additional step to wash bacterial cells prior to adding 5% sucrose was added to remove residual antibiotics and improve consistency of CFU counts.

### Statistical Analysis

To analyze the relationship of multiple *S. mutans* with caries, Fisher’s Exact Test was used. All other statistical analysis was performed using Student’s t-test with statistical significance set to (p<0.05).

## ACKNOWLEDGEMENTS

This research was supported by National Institute of Dental and Craniofacial Research (NIDCR) R01DE016684(N.K. Childers), R01DE022350(H. Wu), T90DE022736(H. Wu), and Diversity Supplement to R01DE022350(S. Momeni).

Special thanks to the Uniontown Study organizers and participants. CLSM imaging and time-lapse movies provided by OHSU Advanced Light Microscopy Core.

